# Exposing the Exposome with Global Metabolomics and Cognitive Computing

**DOI:** 10.1101/145722

**Authors:** Benedikt Warth, Scott Spangler, Mingliang Fang, Caroline H Johnson, Erica M Forsberg, Ana Granados, Richard L Martin, Xavi Domingo, Tao Huan, Duane Rinehart, J Rafael Montenegro-Burke, Brian Hilmers, Aries Aisporna, Linh T Hoang, Winnie Uritboonthai, Paul Benton, Susan D Richardson, Antony J Williams, Gary Siuzdak

## Abstract

Concurrent exposure to a wide variety of xenobiotics and their combined toxic effects can play a pivotal role in health and disease, yet are largely unexplored. Investigating the totality of these exposures, i.e. the *exposome*, and their specific biological effects constitutes a new paradigm for environmental health but still lacks high-throughput, user-friendly technology. We demonstrate the utility of mass spectrometry-based global exposure metabolomics combined with tailored database queries and cognitive computing for comprehensive exposure assessment and the straightforward elucidation of biological effects. The METLIN Exposome database has been redesigned to help identify environmental toxicants, food contaminants and supplements, drugs, and antibiotics as well as their biotransformation products, through its expansion with over 700,000 chemical structures to now include more than 950,000 unique small molecules. More importantly, we demonstrate how the XCMS/METLIN platform now allows for the readout of the biological effect of a toxicant through metabolomic-derived pathway analysis and further, cognitive computing provides a means of assessing the role of a potential toxicant. The presented workflow addresses many of the outstanding methodological challenges current exposome research is facing and will serve to gain a deeper understanding of the impact of environmental exposures and combinatory toxic effects on human health.

---

The concept of the so-called ‘exposome’, the totality of environmental exposures that an individual experiences over their lifetime, was defined over a decade ago to draw attention to the pressing need for innovative methodological developments in exposure assessment^1,2^. It is well-known that environmental exposures play a pivotal role in the etiology of human disease and that the estimated hundreds of thousands of environmental chemicals can be linked to about 80-85% of human disease^3^. However, little is known about the complex interplay between environmental exposures, genetic susceptibility and potential health outcomes. While we are constantly facing a multitude of exposures from the environment, diet, behavior and endogenous processes^4^, exposure and risk assessment is generally still based on the evaluation of single substances or substance classes^5,6^. However, recent initiatives in systems toxicology, and the comprehensive description of toxicological interactions within a living system, are aiming towards the investigation of holistic relationships in a biological system by gaining mechanistic knowledge through combining advanced analytical and computational tools^7,8^.

Major challenges in exposome research include (i) typically very low levels of environmental chemicals or their biotransformation products in the biological samples, and varying concentration ranges between classes of toxicants by orders of magnitude, (ii) a lack of comprehensive analytical workflows covering multiple toxicant classes simultaneously, (iii) compound databases which are almost certainly incomplete or offer only partial coverage and lack MS/MS spectra to facilitate structural identification, (iv) lack of an automated data processing platform and (v) difficulties in correlating biological/health effects with toxicant exposure. As a result, novel analytical methods that are broad in their analyte spectrum, and highly sensitive and specific, are urgently required to better characterize environmental chemicals and their metabolites in biological matrices. High-resolution mass spectrometry (HR-MS) has been proposed to have the potential of being a key technology in exposomics^9,10^. Initial workflows have also proven their applicability for specific applications^11-15^, though untargeted metabolomics has not yet been applied to multi-class exposure assessment in biological fluids to the best of our knowledge. Detailed databases, such as T3DB currently housing approximately 3,700 toxicants^16^, or the Hazardous Substances Data Bank (HSDB; https://toxnet.nlm.nih.gov/newtoxnet/hsdb.htm) with data on more than 5,000 toxicants, have been launched in recent years and constitute very valuable resources.

Beyond detecting toxicants, untargeted metabolomics can play a unique role in exposure research by assessing metabolic changes resulting from the exposure, and hence identifying affected pathways and mechanisms of action. This untargeted approach contrasts to traditional exposure assessment that typically employs targeted analytical methods for specific toxicant classes (Figure 1).

**Figure 1.**
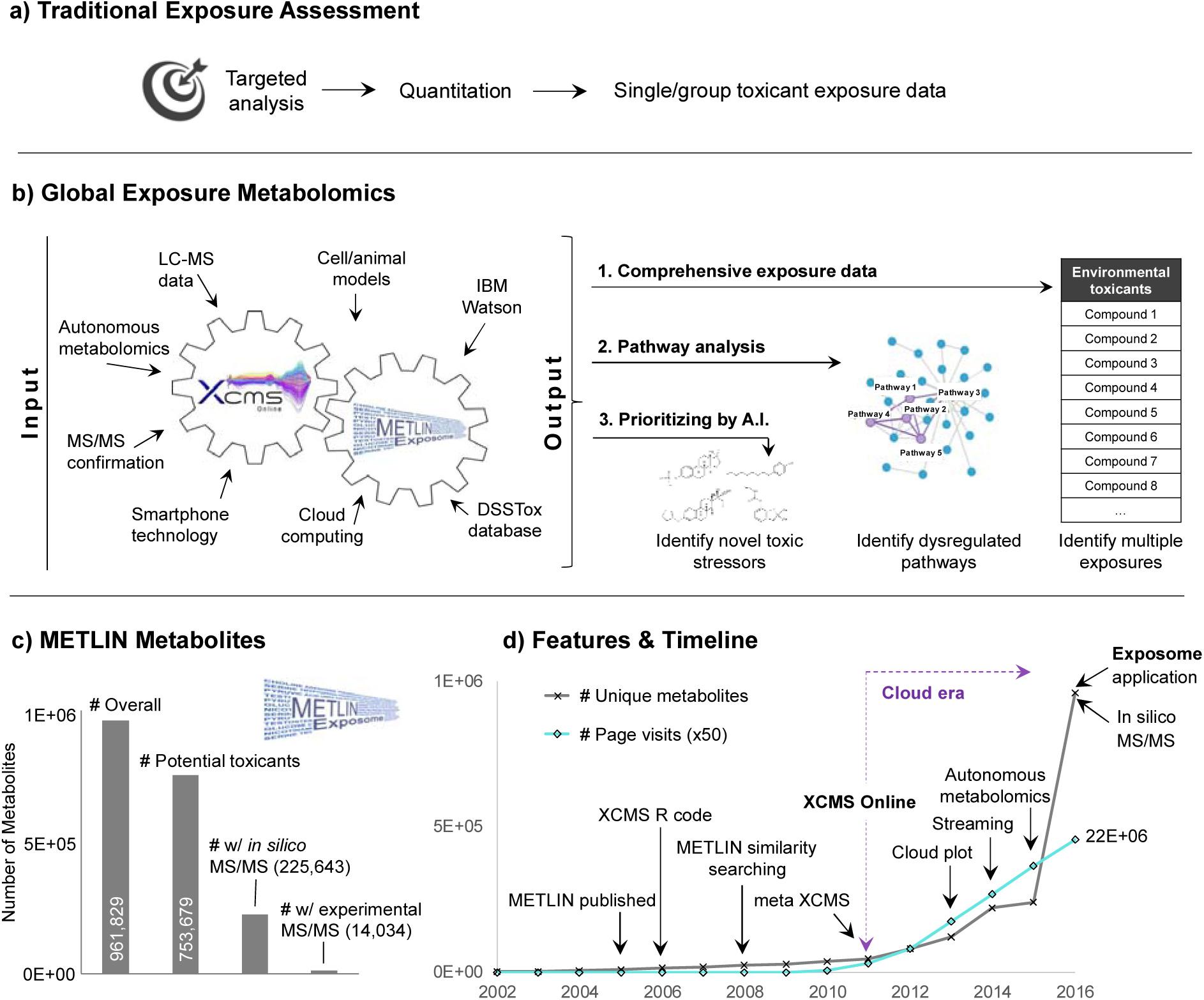
XCMS Online/METLIN platform for environmental and toxicological research. This is a cloud-based, intuitive and user-friendly workflow which enables both, a comprehensive screening of environmental toxicants and a direct biological read-out of affected pathways. In contrast to traditional exposure assessment (a) multiple exposures can be screened simultaneously and used for identifying priorities (b). With >950,000 unique substances, METLIN is the largest metabolite data repository and now includes >750.000 potential toxicants extracted from the DSSTox database (c). MS/MS data for structural identification is available for a large share of molecules. (d) Besides the large number of molecules (black line) and abundant spectral information, the increasing popularity of the METLIN database is explained by the continuous integration of new functionalities which are all well-established in the metabolomics field and are expected to serve exposome-wide studies. Since 2010 more than 22 million METLIN pages were visited by users from 120+ countries as illustrated by the number of cumulative page visits in (d, turquoise line).

Here, we present an untargeted metabolomics workflow that employs HR-MS and freely available XCMS/METLIN cloud-based data processing to examine thousands of metabolic features originating from endogenous metabolites, toxicants, other xenobiotics and their biotransformation products. This concept addresses several of the challenges currently hampering exposome research, and specifically provides a solution to the two key tasks of (1) assessing multiple exposures simultaneously and (2) evaluating their biological effect. The first task may be achieved via a major expansion of METLIN by more than 700,000 chemicals, many of them classed as potential xenobiotics and toxicants, including experimentally observed or *in silico* generated MS/MS spectra for a large share of the library. The second objective is accomplished by implementing a ‘one-click pathway’ analysis technology (Huan et al. *accepted manuscript*) within XCMS Online that highlights significantly modified pathways as a result of exposure. Finally, we demonstrate in three proof-of-principle experiments that exposure to chemicals with endocrine disrupting potential can be determined in human serum and urine at low nanomolar (nM) concentrations, that these chemicals significantly alter cellular metabolism, and that cognitive computing is a promising tool for the purpose of prioritization.

## Results & Discussion

### The METLIN Exposome Database

The identification of endogenous metabolites, as well as of xenobiotics and their biotransformation products, remains a key challenge in both metabolome and exposome research. State-of-the-art HR-MS, which mainly employs quadrupole time-of-flight (Q-TOF) and Orbitrap™ instruments, can enable the annotation of thousands of metabolic features in a single sample. These features may be assigned to specific molecules based on their accurate mass and MS/MS matching using tailored databases such as the Human Metabolome Database (HMDB)^17^, METLIN^18,19^ or others^20^.

To create a comprehensive exposome database, we expanded METLIN by >700,000 chemicals including toxicants and xenobiotics from the United States Environmental Protection Agency’s (EPA) ‘Distributed Structure-Searchable Toxicity (DSSTox)’ database^21^, the dataset underlying the EPA’s CompTox Chemistry Dashboard^22^. Combined, METLIN now covers more than 950,000 unique small molecules ranging from exogenous drugs/metabolites, toxicants, contaminants, lipids, steroids, small peptides, carbohydrates and central carbon metabolites. The DSSTox database includes a manual curation process and for many of the chemicals there is high confidence in the chemical abstract service (CAS) registry number-name-structure with associated chemical structure files for the larger environmental health community. The content of the DSSTox database includes high-production volume chemicals, drugs, disinfection by-products, and chemicals associated with endocrine-related endpoints, carcinogenicity, and aquatic toxicity^21^. While there is limited to no toxicity data available for many of the chemicals due to the very broad scope of the database, a majority of chemicals for which *in vivo* animal data is available were incorporated^13^.

For METLIN, over 16,000 compounds have been individually analyzed to obtain HR tandem mass spectrometric data (MS/MS) at different collision energies. Other databases such as the HMDB^17^ or Massbank^23^ exist, however, these databases are typically represented by a heterogeneous amalgam of data. METLIN includes a substantial share of toxicants (>500) with MS/MS data, however, it is a continuous and ongoing effort to collect HR MS/MS data for additional toxicants. In addition to these experimental spectra, more than 200,000 molecules have MS/MS data generated *in silico* taking advantage of machine learning based on the significant foundation of experimentally acquired MS/MS data generated at different collisional energies (*unpublished)*. This makes METLIN the largest metabolite data repository currently available (Figure 1).

### XCMS Online/METLIN Exposome query functions

XCMS was originally developed as a command line driven metabolomics processing tool in 2006 and evolved into an intuitive, cloud-based platform referred to as XCMS Online in 2011^24^. To enhance the accessibility of XCMS Online and METLIN for exposome-scale investigations, and to tailor the platform according to the needs of the community, several search functions have been implemented. For example, filtering for either specific toxicants or all toxicants within a given dataset can be performed (see Supplementary Figure S1 for screenshots):

1. **Quick compound search:** Using this filter in the ‘results table’, only features containing a hit for the specified keyword (e.g. ‘triclosan’) within the METLIN database, and matching a specified mass accuracy (e.g. 5 ppm), are displayed. The additional mass filter included on the keyword search helps to reduce the number of presented toxicants to only those corresponding to the possible ion species. The extracted features may then be evaluated manually on their plausibility taking isotopic information, adduct formation and retention behavior into account.
2. **Toxicants unknown search:** Using this filter, exclusively metabolic features containing hits for toxicants are presented in the results table. This enables the targeted screening for unknown compounds in the samples including man-made toxicants, natural toxicants, drugs and other metabolites of non-endogenous origin in an unbiased manner. Also, for this functionality, the extracted features may then be evaluated and validated manually. Supplementary Table S1 summarizes the xenobiotics detected in pooled human serum and urine samples by applying this novel multi-class screening approach.
3. **METLIN toxicant search:** Within the METLIN database it is now possible to specifically include or exclude toxicants from a search query. This function serves to find solitary toxicant entries associated with a mass, name, chemical formula, KEGG^25^ or CAS Registry Number.
4. **METLIN MS/MS search:** Users can choose to include only toxicant entries containing experimental or *in silico* MS/MS data facilitating structural information.

### Global Untargeted Screening

To evaluate the ability of the XCMS Online/METLIN exposome workflow to screen for xenobiotics in an unbiased manner, we analyzed one of each pooled human serum and plasma samples. Many different exogenous molecules were detected including drugs, antibiotics, dietary compounds, food and environmental contaminants. A list containing detailed information regarding mass accuracy, ion species, MS/MS spectra, and classification is presented in Supplementary Table S1.

It is important to mention that, despite the number of toxicants and drugs detected in this proof-of-principle study, the levels of many environmental toxicants are still too low to be observed by the established untargeted screening approach, and some may not be ionized by electrospray ionization. However, this might be overcome in part by optimizing sample cleanup and enrichment procedures in the future. To evaluate the analytical performance of the established method, spiking experiments were performed. Three well-known xenobiotics of different chemical classes were spiked into a pooled serum sample at a low (5 nM) and high (500 nM) concentration. This includes the isoflavone genistein (GEN: https://comptox.epa.gov/dashboard/DTXSID5022308) a common food supplement, the mycotoxin zearalenone (ZEN: https://comptox.epa.gov/dashboard/DTXSID0021460), a frequent contaminant of grain-based food and the polychloro phenoxy phenol triclosan (TCS: https://comptox.epa.gov/dashboard/DTXSID5032498), an antibacterial agent used in personal care products. Method detection limits (MDL) of the model compounds, which are all controversially discussed in the context of their potential endocrine disrupting properties, could be estimated based on their signal-to-noise ratios. As illustrated in Supplementary Figure S2 the method was able to detect all three compounds at concentrations which constitute a realistic baseline scenario in epidemiological studies. Under the chosen chromatographic and electrospray ionization conditions, the MDLs of GEN, ZEN and TCS were estimated to be approximately 2, 4 and 38 nM, respectively (corresponding to 0.5, 1.3 and 11 ng/mL). Although the method did not employ any specific cleanup, we were able to detect GEN even in the non-spiked serum sample, while its absence was confirmed in the solvent blanks. Based on a relative comparison with the samples spiked at a concentration of 5 nM (1.4 ng/mL) the background concentration was estimated to be approximately 6 nM, a concentration range reported previously in healthy humans^26^. However, higher exposures up to 378 nM have been reported as mean plasma concentration in women from the United Kingdom (UK) with high soy intake^27^ and suggest that our methodology is well suited for detecting low and high exposures of GEN. The same holds true for ZEN, for which no reliable data on serum concentrations are available, and exposures in the range of the tolerable daily intake (TDI) resulted in urinary concentrations in the low nM range^28^, and TCS for which relevant concentrations are typically in the nM range^29,30^. This confirms that the methodology is fit-for-purpose for the investigated model xenobiotics. However, further enhancements in sensitivity might be achieved through the application of more tailored sample preparation protocols or more sensitive mass spectrometers.

A further asset of the METLIN exposome database is the inclusion of a significant number of mammalian biotransformation products. This includes, for example glucuronides (726 total including 654 with associated MS/MS information) and sulfate-conjugates (1989 total with 366 containing MS/MS data) but also abundant phase I products (hydroxylation, methylation, etc.). Consequently, it is possible to directly evaluate the metabolism/toxicokinetics of a toxicant in a specific individual or population sub-group in studies collecting samples at multiple time points, an potentially valuable tool when evaluating for detoxification capacities. A current example of how diverse the metabolism of a xenobiotic may be dependent on species, gender and hormone levels was exemplified by Soukup, et al.^31^ for GEN and other isoflavones. Hence, we additionally investigated the presence of glucuronide and sulfate conjugates in the pooled serum sample. Based on accurate mass and retention behavior we putatively identified a mono- and a di-glucuronide (Table S1). Thereby, we demonstrated the potential of HR-MS to screen for conjugates and other metabolic products for which no reference standards are commercially available to date. As expected, we did not detect the conjugates in urine because these samples were deconjugated utilizing b-glucuronidase/acrylsulfatase (see methods section).

### Toxicological Pathway Analysis

In addition to the METLIN database expansion, perhaps the most significant advantage of the presented platform is its ability to go beyond exposure assessment by inherently depicting affected pathways in human samples or cell-based models. Taking advantage of the global metabolomics approach, hundreds of endogenous metabolites are measured simultaneously with the analysis of the toxicants screened for exposure assessment. Utilizing the newly developed ‘pathway cloud plot’ feature within XCMS Online (Huan et al. *accepted manuscript*) allows for a straightforward readout of affected pathways, thus enabling an investigation of the underlying mode of action. A major issue in current exposome research is deciphering which toxicant has had the metabolic/biological effect in an epidemiological study. We propose initial dosing in cell or mouse models with individual toxicants and evaluating their effect, followed by gradually adding more substances to see the combined effects, might be one future experimental solution to this challenge. A similar approach was used in a study that investigated reproductive effects of regulated disinfection by-product mixtures (trihalomethanes and haloacetic acids) on the offspring of rats that were gestationally exposed^32^. We believe the presented workflow offers researchers a unique resource to evaluate single or multiple exposures and receive a direct readout of the altered metabolic pathways in a straightforward one-step procedure.

To demonstrate the potential of this novel intrinsic pathway analysis to evaluate a toxicant’s mode of action, we incubated MCF-7 breast cancer cells with a low concentration (0.1 μM) of the xenoestrogen ZEN in another proof-of-principle experiment. The metabolomics cloud plot in Figure 2a displays all metabolic features significantly dysregulated as a result of the perturbation while the newly developed pathway cloud plot (Figure 2b) links these features to specific pathways (Huan et al. *accepted manuscript*). Thus, it facilitates direct information on the modulated pathways of any xenobiotic or mixture which is a unique, integrated advantage of the applied workflow and may be a valuable tool in systems toxicology. Another major advantage of this pathway cloud plot is the fact that the manual annotation and interpretation of MS/MS spectra for identification purpose are avoided. This is caused by the algorithm based on the *mummichog* concept, predicting pathways directly from spectral feature tables without the need for *a priori* identification of metabolites^33^. Consequently, this results in a significantly faster data evaluation process which is crucial when evaluating large scale epidemiology studies or cell models of toxicant combinations which both typically involve high sample numbers.

**Figure 2.**
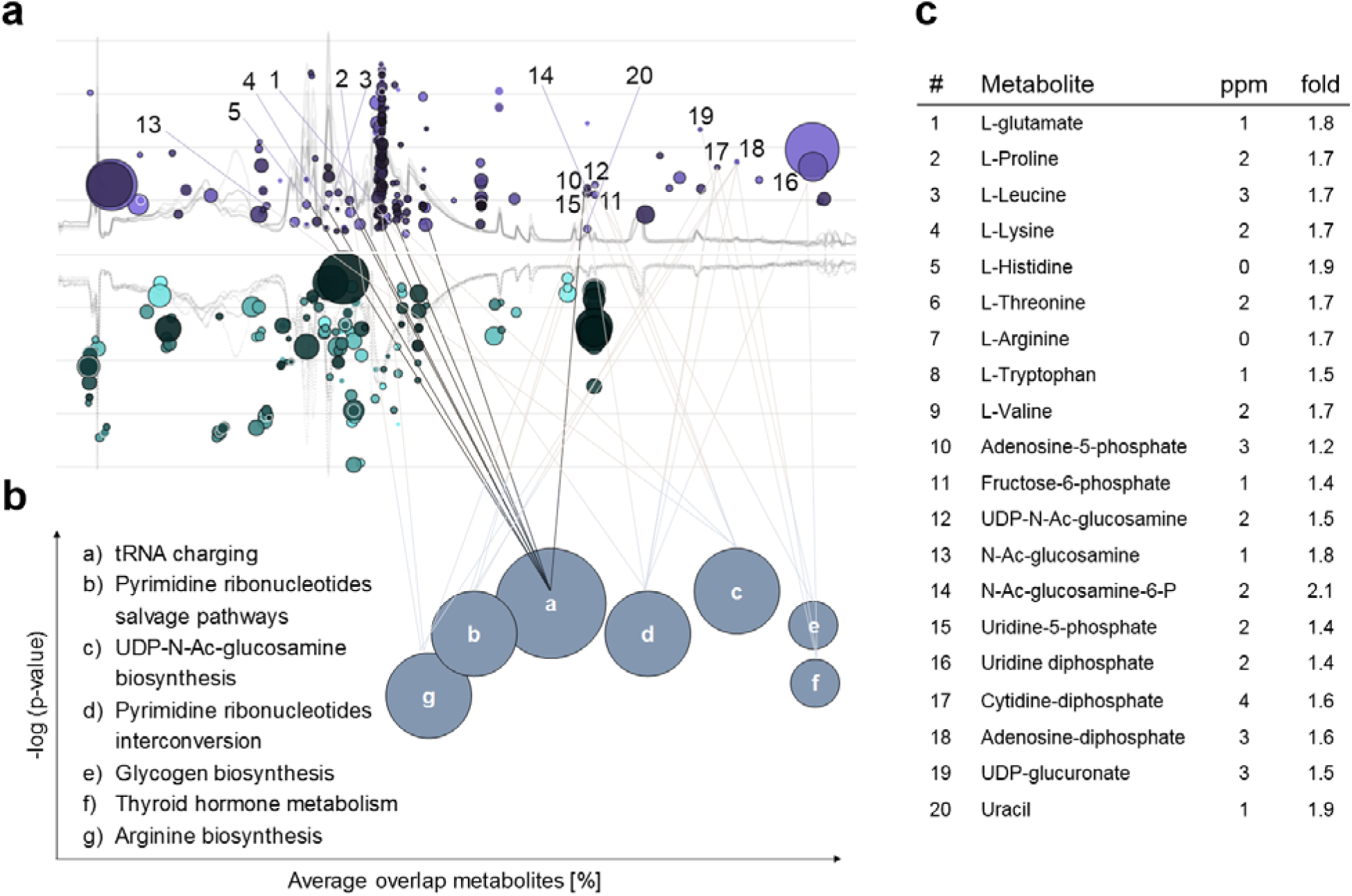
Metabolic pathway analysis aiming towards systems toxicology in a cell model. In the metabolomics cloud plot (a) metabolic features significantly altered as a result of 48 h xenoestrogen exposure (0.1 μM zearalenone; ZEN) in the MCF-7 breast cancer cell line compared to control treatment are illustrated. The color intensity of a bubble represents the *p*-value (all < 0.05; the more intense the lower) whereas purple and turquoise indicate up- or down regulation following exposure, respectively. The radius represents the fold change of a specific metabolic feature. The schematic pathway cloud plot in (b) maps perturbations in cellular metabolism caused by the xenoestrogen and translates the individual metabolic features reported in (a) to the systems biology/toxicology level. Pathways were embedded into XCMS Online from the BioCyc database and are mapped using the *mummichog* algorithm. In (c) selected underlying endogenous key metabolites and their fold change are displayed. All data processing and analysis was performed using XCMS Online.

In our pilot experiment, the xenoestrogen ZEN clearly led to increased abundances of amino acids, nucleosides and nucleotides, while glycerol-phosphate and hypoxanthine were down-regulated. The pathways associated with these changes were automatically detected by the new XCMS Online functionality and manually verified (Figure 2c) using reference standards and MS/MS spectra. While no global metabolomics study was carried out on ZEN exposed MCF-7 cells to date, our results are supported by a study carried out in murine macrophage ANA-1 cells in which a potent effect on amino acid metabolism was discovered as well^34^. The next phase of this effort is to investigate epidemiological studies with numerous confounding factors and co-exposures. However, it is likely that specific exposure patterns (e.g. high overall exposure to EDCs) might be correlated to affected pathways in large cohort studies.

### Exploring cognitive computing utilizing Watson

Given the significant number of candidates presented in this study and added to METLIN, we investigated artificial intelligence to prioritize findings. Artificial intelligence/machine learning was employed for accelerated data mining with IBM Watson^®^. Here, we used Watson to screen 15,000 toxicant candidates from the EPA’s DSSTox list, for potentially new EDCs. A total of 15,054 toxicant candidates were entered as queries to Watson, of these, 9,502 were identified as unambiguous chemical structures. Using a training set of 30 known estrogen receptor (ER) agonists selected from the EPA’s Endocrine Disruptor Screening Program (EDSP) and a semi-supervised learning approach (graph diffusion) for classification, 991 candidates could be ranked (Supplementary Table S2). 20 known ER agonists were utilized for validation purposes and integrated into the candidate set, 11 of those were recognized by Watson. Thereof, a fraction of 92% was within the top 10% rank order (Table S2). Besides the training set, Watson was able to identify a number of other EDCs described in the literature which were among the 15,000 toxicant candidates.

In these analyses over 313,000 Medline^®^ abstracts were examined where chemical names are input as a query, and Watson then attempts to interpret each input name as an unambiguous chemical structure (see Online Methods for technical details), and for each structure will then return the documents associated with that compound. Name formats understood by Watson include generic, brand, and common names, as well as structure representations such as SMILES [http://www.daylight.com/dayhtml/doc/theory/theory.smiles.html] and InChl [http://www.inchi-trust.org/]. Searching Watson for a particular input name will therefore retrieve all documents where the corresponding chemical was mentioned, comprising documents where the text includes an exact match to the specific input, as well as documents containing any other synonymous chemical representation. A reader capable of absorbing 10 papers per day would need nearly 100 years to go through this potentially relevant literature, an unrealistic feat. Instead, Watson^35^ mines text so as to create a model for each metabolite that represents all the terms present in the abstracts of the papers that specifically mention the named metabolite.

We believe that this example demonstrates the clear predictive capability of Watson in finding new potential EDCs similar to the training set. More importantly, cognitive computing is not limited to a specific mode of mechanism and may be extended to other toxicant classes such as carcinogens or genotoxic compounds. Therefore, the evaluated machine learning strategy provides a valuable resource for future identification of suspects and literature based priority ranking. This holds the potential to screen tens to hundreds of thousands of chemicals in some hours/days which would not be possible manually and potentially opens up new and unexpected discoveries. We believe this approach is of special value to the exposomics field dealing with extensive data sets and will be even more powerful once a second generation of machine learning algorithms are able to answer structure-based queries.

## Conclusion & Outlook

A comprehensive untargeted screening of xenobiotics and their biological effects was developed and its applicability examined in human biofluids and a cell-based model. This approach involves HR-MS analysis and freely available cloud-based data processing and evaluation at the systems toxicology level. It is expected to support large-scale epidemiological studies for initial toxicant screening, hypothesis generation and the elucidation of underlying mechanisms of toxicity. Furthermore, it may be readily adopted to other fields such as food, feed, water, air and soil analysis to serve a multitude of scientific communities. This workflow enables, for the first time, the combination of holistic, untargeted exposure assessment with changes in endogenous metabolism for deciphering specifically affected pathways. Moreover, we demonstrated the vast potential of artificial intelligence to prioritize findings obtained from global metabolomics experiments based on toxicological mechanisms within the framework of exposome-scale studies. Good data sharing practices and smart linking of online resources, as demonstrated here with XCMS/METLIN, will be important to further integrate data evaluation in exposome research. We believe the presented technology platform, and the three presented proof of principle experiments involving potentially endocrine disrupting chemicals, have the potential to influence how we address the challenges in this new era of exposomics and systems toxicology evolution.

## Methods

### Chemicals & Reagents

Acetonitrile (ACN; Fisher Scientific, Pittsburgh, PA), methanol (MeOH; Honeywell, Morris Plains, NJ) and water (J.T. Baker, Center Valley, PA) were all LC-MS grade. Dimethyl sulphoxide (DMSO) (≥ 99,5%), GEN, ZEN and TRI as well as β-glucuronidase/acrylsulfatase were purchased from Sigma-Aldrich (St. Louis, USA). Cell culture media and supplements were purchased from ATCC (Manassas, Virginia) and Fisher Scientific (Pittsburgh, PA).

### Human samples

A pooled human serum (male AB plasma, USA origin, sterile-filtered) for spiking experiments was purchased from Sigma-Aldrich (St. Louis, USA), aliquoted on ice and immediately stored at −80°C. A pooled human urine sample was prepared by mixing first morning samples of eight volunteers (both gender, age 28-50) which were immediately frozen at −20°C after sampling. No information on dietary or lifestyle habits and medical status was collected to allow for unbiased screening of toxicants, dietary contaminants and drugs. Informed written consent was obtained from all participants. This study was approved by the Scripps Institutional Review Board (IRB NUMBER: IRB-16-6919).

### Sample preparation

Human serum samples (200 μL) were extracted with 800 μL of cold MeOH:ACN (1:1, v/v), vortexed for 30 s and sonicated for 10 min. To precipitate proteins, samples were incubated for 1 h at −20 °C, followed by 15 min centrifugation at 13.000 rpm and 4°C. The supernatant was evaporated to dryness in a vacuum concentrator (Labconco, Kansas City, MO) and the dried extracts reconstituted in 100 μL of ACN:H2O (1:1, v/v). Following sonication for 10 min, and centrifugation (15 min) at 13.000 rpm and 4°C, the supernatants were transferred to LC vials and stored at −80 °C prior to analysis. Urine samples (0.5 mL each) were centrifuged for 10 min at 13,000 rpm and 4°C and the supernatant transferred to a fresh tube. Then, the urine samples were diluted 1:1 with PBS and enzymatically treated with b-glucuronidase/acrylsulfatase for 18 h at 37°C. Then, the treated urine samples were diluted with 1 mL ACN, put to −20°C for 1h and centrifuged (10 min, 13,000 rpm, 4°C). The supernatants were evaporated to dryness utilizing a vacuum concentrator and reconstituted in either 0.2 mL of ACN:H2O (1:1, v/v), centrifuged and transferred to an LC vial (overall enrichment factor 2.5). MDLs were estimated based on a signal to noise ratio of 3:1. Spiking experiments were done by adding a mixed stock solution of the compounds to the pooled sample before sample preparation. Since GEN was present in the samples, standard addition was performed.

### Cell culture

MCF-7 breast cancer cells (ATCC, Manassas, VA) were cultured in Dulbecco’s Modified Eagle Medium F-12 Nutrient Mixture (Gibco-Life Tech, Grand Island, NY) supplemented with 10% fetal bovine serum (Sigma, St. Louis, MO) and penicillin-streptomycin (50 U/mL, Gibco-Life Tech, Grand Island, NY) at 37°C and 5% CO_2_. Prior to the experiments the cells were split routinely and maintained in T-75 flasks at a confluence between 75-85%. For sub-culturing the cells were washed with PBS, detached with trypsin/EDTA and re-suspended in fresh culture medium. For the experiments cells were seeded in 6- well plates. After 48 h cells were either incubated with common medium (control) or 0.5 μM ZEN. All experiments were performed in triplicate.

To extract the cell lysates, the 6-well plates were put on ice and the medium in the wells was removed using a vacuum pump. Cells were washed with 1 mL of ice cold PBS twice. Then, 1 mL ice cold quenching solution (MeOH:ACN:H2O (2:2:1, v/v) was added and the cells were detached using a cell scraper and the cell suspension was transferred to Eppendorf tubes. Samples were vortexed followed by three cycles of shock-freezing in liquid nitrogen and subsequent thawing at room temperature and sonication for 10 min. To precipitate proteins samples were incubated for 1 h at −20 °C, followed by 15 min centrifugation at 13,000 rpm and 4°C. The supernatant was evaporated to dryness in a vacuum concentrator (Labconco) and the dried extracts reconstituted in ACN:H2O (1:1, v/v) according to their protein content as determined by the BCA assay. Following sonication for 10 min, and centrifugation (15 min) at 13,000 rpm and 4°C the supernatants were transferred to LC vials and stored at −80 °C prior to analysis.

### LC-MS/MS instrumentation

Serum, urine and cell extract analyses were performed using a high performance liquid chromatography (HPLC) system (1200 series, Agilent Technologies) coupled to a Bruker Impact II quadrupole time-of-flight (Q-TOF) mass spectrometer (Bruker Daltonics, Billerica, MA). Serum and urine samples were injected (4 μL) onto an Atlantis T3, 3 μm, 150 mm × 1.0 mm I.D. column (Waters, MA) for reversed phase analysis in ESI positive and negative mode. The mobile phase was A = 0.1% formic acid in water and B = 0.1% formic acid in acetonitrile. The linear gradient elution from 5 % B (0-5 min) to 100 % B (40-45 min) following 7 min of column re-equilibration at 100% B was applied at a flow rate of 60 μL/min. ESI source conditions were set as follows: gas temperature 220°C, drying gas (nitrogen) 6 L/min, nebulizer 1.6 bar, capillary voltage 4500 V. The instrument was set to acquire over a *m/z* range from 50-1000 with the MS acquisition rate of 2 Hz. Cell extracts were injected (4 μL) onto a Luna aminopropyl, 3 μm, 150 mm × 1.0 mm I.D. column (Phenomenex, Torrance, CA) for HILIC analysis in ESI negative mode. HILIC was chosen to analyze predominantly central carbon metabolites which typically retain better by HILIC than revered phase columns. The mobile phase was A 20 mM ammonium acetate and 40 mM ammonium hydroxide in 95% water and 5 % acetonitrile and B 95% acetonitrile, 5 % water. The linear gradient elution from 100 % B (0-5 min) to 100 % A (50-60 min) was applied in HILIC at a flow rate of 50 μL/min. To ensure column re-equilibration and maintain reproducibility a 12 min post-run was applied. For the acquisition of MS/MS spectra of selected precursors the default isolation width was set to 0.5 Da with MS and MS/MS acquisition rates of 4 Hz. The collision energy was set to 20-50 eV.

### XCMS Online data processing (global metabolomics)

Data was processed using XCMS Online^36^ with an α=0.05. Data was displayed as *a* feature table and plotted as cloud plot for both, the metabolomic and pathway cloud plots. These contained *m/z* and retention time information (where applicable), integrated intensities, observed fold changes across the sample groups, and statistical significance for each sample. The web server log was used to track the number of METLIN page visits since 2010. Usage statistics in Figure 1 (c) are reported as cumulative pages since 2010.

### Cognitive computing

50 known ER agonists were selected from the EPA’s Endocrine Disruptor Screening Program (EDSP; Supplementary Table S2) and randomly divided into a training set consisting of 30 compounds and a validation set of 20 molecules. The latter ER agonists were integrated in the candidate set which consisted of a randomly selected subset of 15,000 chemicals from the EPA’s DSSTox list.

The Watson system combines model-based and rule-based natural language processing technologies to identify chemical names as they occur in natural language text. It also comprises a cheminformatics post-processing component which interprets these names as unambiguous chemical structures. An example of the model-based system for identifying systematic chemical names within text is the use of machine learning to train a system to recognize systematic, IUPAC-style chemical names^37^. By contrast, an example of the rule-based approach is to extract known chemical names such as generic and common names from text based upon a dictionary of reliable chemical terms, assembled from an assortment of external expert data sources. These methods are complementary, and together constitute a *hybrid* approach to natural language processing, which is intended to mimic human expert parsing of language by examining both the *content* and the *context* of phrases in text. For example, a model-based approach can be particularly performant for identifying chemical names in text whose content conforms to a systematic pattern (i.e. the combination of alphanumeric characters and symbols that comprise an IUPAC name such as “1,3,7-Trimethylpurine-2,6-dione”), while a rule-based approach utilizing a dictionary can be particularly performant for identifying generic or common names in text based on their content (i.e. names that may not conform to any systematic pattern, such as “caffeine”). Similarly, both model-based and rule-based methods can be performant in the extraction of chemical names based upon their context, e.g. based upon the semantic decomposition of a sentence.

Whenever a chemical name is discovered in text or entered as a query, Watson attempts to generate an unambiguous chemical structure for that name. Indeed, each chemical name identified in text must also pass this automated analysis step, and successfully be converted into a chemical structure. The purpose of this conversion is two-fold: to consolidate synonymous chemical names under a single canonical representation, for example to recognize that the name “caffeine” indeed refers to the same compound as “1,3,7-Trimethylpurine-2,6-dione”, and also to eliminate names that cannot be understood as unambiguous, single chemical structures, for example phrases detected in the natural language processing step that are not single chemicals (but rather chemical classes, non-chemicals, and so on). A cheminformatics component within the Watson system is responsible for this conversion, and utilizes both third-party cheminformatics software tools and in-house approaches to interpret chemical names, including pre-processing of the names and post-processing of the results to ensure high-quality chemical structure generation. For the purposes of indexing within the Watson system, each computed chemical structure is represented unambiguously with a canonical SMILES string, i.e. a textual notation format for chemical structures based upon interpreting the chemical graph as a spanning tree [http://www.daylight.com/dayhtml/doc/theory/theory.smiles.html].

Watson collected and labeled all available abstracts to be mined using queries against a text index of all Medline abstracts. It submitted the queries and downloaded all abstracts that match each metabolite up to query size. Next, a numeric representation that encapsulates all that is known about each metabolite relative to every other metabolite was created in a vector space model. Each document is represented by a vector of weighted frequencies of its features (words and phrases)^38^. Various standard stop words are excluded from the document in a first pass, resulting in a rank of N words. Watson finds approximately 20,000 words is a suitable approximation for N. As a second pass, phrases are considered where no ‘stop’ word occurs with two words in order, this is the features space. Finally, a third pass over the document indexes the feature frequency. Various methods of weighting terms occurrences were tested with a Term Frequency-Inverse Document Frequency weighting (TF-IDF)^38^ yielding the best overall prediction accuracy.

To prioritize metabolites for future experiments, graph diffusion^39^ was used as a ranking scheme. Here, is our function that we need to solve. Our y term is the fixed binary vector to tables such that y represents if a metabolite I has ER ligand properties. To solve *f* we can minimize the sum losses (*f* – *y*)*^T^* & the smoothing term (*f* – *y*) our second term. The Laplacian matrix is our representation of the metabolite network, defined by *L* = *D* – *A*. Here A is our adjency matrix and D is our Degree matrix. If we set our diffusion coefficient *μ*, to the inverse of the Laplacian norm we can generate a closed form equation eq2^40^:

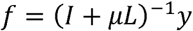

In order to identify new metabolites we then look at the new labels *f* where the labels with the largest increase will be our targets. To validate the obtained ranking a *Leave-One-Out* cross validation approach was utilized, doing one run for each training entity by leaving that entity out of the training set and evaluating how it ranks as a candidate. A ROC curve for the validation and recall is shown in Supplementary Figure S3.

## Data Availability

The data sets presented in this study are publicly available via XCMS Public (https://xcmsonline.scripps.edu) under the job numbers #1137610, #1137611, #1129233, and #1136439 (will be made available once accepted for publication).

## Author contributions

B.W., M.F., and G.S. designed the research idea and experiments. B.W., A.G., R.M., and S.S. performed the experiments. B.W., M.F., C.H.J., A.G., X.D., T.H., S.D.R., and G.S. analyzed and evaluated the data. B.W., D.R., A.A., P.P.B., and G.S. contributed to the development, implementation, and optimization of the METLIN Exposome database and the new XCMS Online functionalities. A.J.W. provided the DSSTox database and helped in experimental design. All authors contributed to manuscript writing.

## Acknowledgements

The authors would like to thank the following colleagues for valuable discussions and/or skillful technical assistance: Minerva Tran (TSRI), Bill Webb (TSRI), Elin M Ulrich (EPA), Jon R Sobus (EPA). We greatly acknowledge the Austrian Science Fund (FWF) for supporting the first author with an Erwin Schrödinger fellowship (J-3808), the National Institutes of Health grants RO1 GMFI4368 and PO1 A1043376-02S1 and Ecosystems and Networks Integrated with Genes and Molecular Assemblies (http://enigma.lbl.gov), a Scientific Focus Area Program at Lawrence Berkeley Laboratory for the U.S. Department of Energy, Office of Science, Office of Biological and Environmental Research under contract number DE-AC02- 05CH11231. The views expressed in this paper are those of the authors and do not necessarily reflect the views or policies of the US Environmental Protection Agency. Mention of trade names or commercial products does not constitute endorsement or recommendation for use.

